# High-resolution map of plastid encoded polymerase binding patterns demonstrates a major role of transcription in chloroplast gene expression

**DOI:** 10.1101/2022.01.18.476797

**Authors:** V. Miguel Palomar, Sarah Jaksich, Sho Fujii, Jan Kuciński, Andrzej T. Wierzbicki

## Abstract

Plastids are endosymbiotic organelles containing their own genomes, which are transcribed by two types of RNA polymerases. One of those enzymes is a bacterial-type, multi-subunit polymerase encoded by the plastid genome. The plastid encoded RNA polymerase (PEP) is required for efficient expression of genes encoding proteins involved in photosynthesis. Despite the importance of PEP, its DNA binding locations have not been studied on the genome-wide scale at high resolution. We established a highly specific approach to detect the genome-wide pattern of PEP binding to chloroplasts DNA using ptChIP-seq. We found that in mature *Arabidopsis thaliana* chloroplasts, PEP has a complex DNA binding pattern with preferential association at genes encoding rRNA, tRNA, and a subset of photosynthetic proteins. Sigma factors SIG2 and SIG6 strongly impact PEP binding to a subset of tRNA genes and have more moderate effects on PEP binding throughout the rest of the genome. PEP binding is commonly enriched on gene promoters, around transcription start sites. Finally, the levels of PEP binding to DNA are correlated with the levels of RNA accumulation, which allowed estimating the quantitative contribution of transcription to RNA accumulation.

## INTRODUCTION

Plastids are endosymbiotic organelles, which contain their own genomes derived from a cyanobacterial ancestor. Plastid genomes are relatively small, containing between 120 and 160 kb of DNA and encoding typically between 100 and 120 genes (Bock, 2007). The Arabidopsis plastid genome encodes 120 genes in 154,478 bp of DNA (Sato et al., 1999). Most plastid encoded proteins and non-coding RNAs are components of gene expression machinery or photosynthetic enzyme complexes (Bock, 2007). The remainder of the complex plastid proteome is encoded by the nuclear genome and transported into plastids post-translationally (Bock, 2007).

Plastid genomes are transcribed by two types of RNA polymerases. The nuclear-encoded RNA polymerase (NEP) is a phage-type, single-subunit enzyme. NEP transcribes mostly housekeeping genes and is most active early in chloroplast development (Ortelt and Link, 2021; Pfannschmidt et al., 2015). The plastid encoded RNA polymerase (PEP) is a bacterial-type enzyme with four core subunits (α, β, β’, and β’’) encoded by the plastid genome (*rpoA*, *rpoB*, *rpoC1*, and *rpoC2*, respectively). It transcribes mostly genes encoding photosynthetic proteins, such as subunits for photosystems and the Rubisco large subunit (RbcL), and is the predominant RNA polymerase in mature chloroplasts (Ortelt and Link, 2021; Pfannschmidt et al., 2015).

Similar to bacterial RNA polymerase, nuclear-encoded sigma factors (SIG) are required for PEP activity by recruiting PEP to gene promoters (Chi et al., 2015; Lysenko, 2007). Six SIG isoforms have been identified in Arabidopsis. Although they have partially redundant functions, loss of SIG2 and SIG6, but not other sigma factors, broadly decreased the mRNA levels of PEP-transcribed genes and impaired chloroplast development in seedlings, indicating the importance of these two sigma factors in chloroplast biogenesis (Woodson et al., 2013). The major targets of SIG2 and SIG6 are considered to be tRNA coding genes and photosynthesis-related genes, respectively (Ishizaki et al., 2005; Kanamaru et al., 2001). A group of peripheral PEP components, pTAC or PAP proteins, is also important for PEP activity (Pfalz and Pfannschmidt, 2013; Pfannschmidt et al., 2015).

Plastid transcription has been studied using run-on experiments designed to assay specific genes in spinach (Deng et al., 1987), potato (Valkov et al., 2009), barley (Krupinska and Apel, 1989; Melonek et al., 2010), tobacco (Krause et al., 2000; Legen et al., 2002; Shiina et al., 1998) and Arabidopsis (Isono et al., 1997; Tsunoyama et al., 2004). Chromatin immunoprecipitation (ChIP) is another approach that has been used to estimate the patterns of transcription by determining the DNA binding pattern of an RNA polymerase. It has been performed in tobacco using epitope-tagged RpoA and genome-wide detection of DNA using a microarray. This method achieved an average spatial resolution of 716 bp, which limits obtained insights to the scale of individual genes (Finster et al., 2013). In Arabidopsis, PEP binding to DNA has been assayed on a limited number of specific loci (Ding et al., 2019; Hanaoka et al., 2012; Yagi et al., 2012) and the genome-wide pattern of PEP activity remains unknown.

Existing run-on and DNA binding data demonstrated substantial differences in transcription and PEP association with DNA between various plastid genes (Deng et al., 1987; Finster et al., 2013). The impact of PEP activity on changes in gene expression in response to developmental or environmental cues is variable with evidence for gene regulation occurring with or without changes in transcription rates (Isono et al., 1997; Krupinska and Apel, 1989; Shiina et al., 1998). Additionally, gene expression in plastids is strongly influenced by posttranscriptional processes including RNA processing and translation (Barkan, 2011; Stern et al., 2010). The impact of transcription on plastid gene regulation remains only partially understood because existing data inform about PEP transcription on limited numbers of loci or with low resolution. Therefore, the pattern of PEP activity within individual genes or operons is known on just a few loci. Moreover, the relationship between transcription and RNA accumulation is unknown on the genome-wide scale. It is also not known how individual sigma factors contribute to recruiting PEP to specific genes.

We established an improved method to study protein-DNA interactions in plastids and applied it to determine the genome-wide pattern of PEP binding to DNA. We confirmed that PEP has a complex pattern of DNA binding and found that PEP binding is the strongest at rRNA and tRNA genes. Sigma factors SIG2 and SIG6 have dual impacts on PEP binding to specific tRNA genes and to the remainder of the genome. PEP associates with a substantial subset of gene promoters and the levels of PEP binding are correlated with steady-state levels of RNA accumulation. Presented data are available through a publicly available Plastid Genome Visualization Tool (Plavisto) at http://plavisto.mcdb.lsa.umich.edu.

## RESULTS

### Genome-wide detection of PEP binding to chloroplast DNA

To detect interactions between PEP and specific sequences within the plastid genome, we adapted a nuclear ChIP-seq protocol (Rowley et al., 2013) for use with chloroplasts. We refer to this method as plastid ChIP-seq (ptChIP-seq). A critical step of ChIP is crosslinking with formaldehyde, which covalently preserves protein-DNA interactions (Hoffman et al., 2015). The ptChIP-seq protocol was designed to maximize capture of protein-DNA interactions and, unlike most applications in the nuclear genome, uses 4% formaldehyde (Davis et al., 2011; Zaidi et al., 2017). To demonstrate the specificity of ptChIP-seq, we first compared different lengths of time of the crosslinking reaction. For this purpose, we used 14 days old plants expressing HA-tagged pTAC12/HEMERA (Galvão et al., 2012). pTAC12 is one of PEP-associated factors (Pfalz et al., 2006), which binds at least a subset of PEP-transcribed loci (Pfalz et al., 2015). Crosslinking for 4 hours resulted in the highest signal to noise levels, compared to 1h and 16h (Fig. 1A and Fig. S1A). This was especially visible on tRNA and rRNA genes, where ptChIP-seq signal was the strongest (Fig. 1A and Fig. S1A). We obtained similar results performing ptChIP-seq using chloroplasts enriched from Col-0 wild-type plants and a commercially available polyclonal antibody against the β subunit of PEP (RpoB; Fig. 1B, Fig. S1B). No signal was observed in non-crosslinked controls (Fig. 1A and Fig. 1B), which indicates that unlike other related protocols (Barkan, 2009; Newell and Gray, 2010), ptChIP-seq only captures protein-nucleic acid interactions that have been preserved by crosslinking.

**Figure 1.**
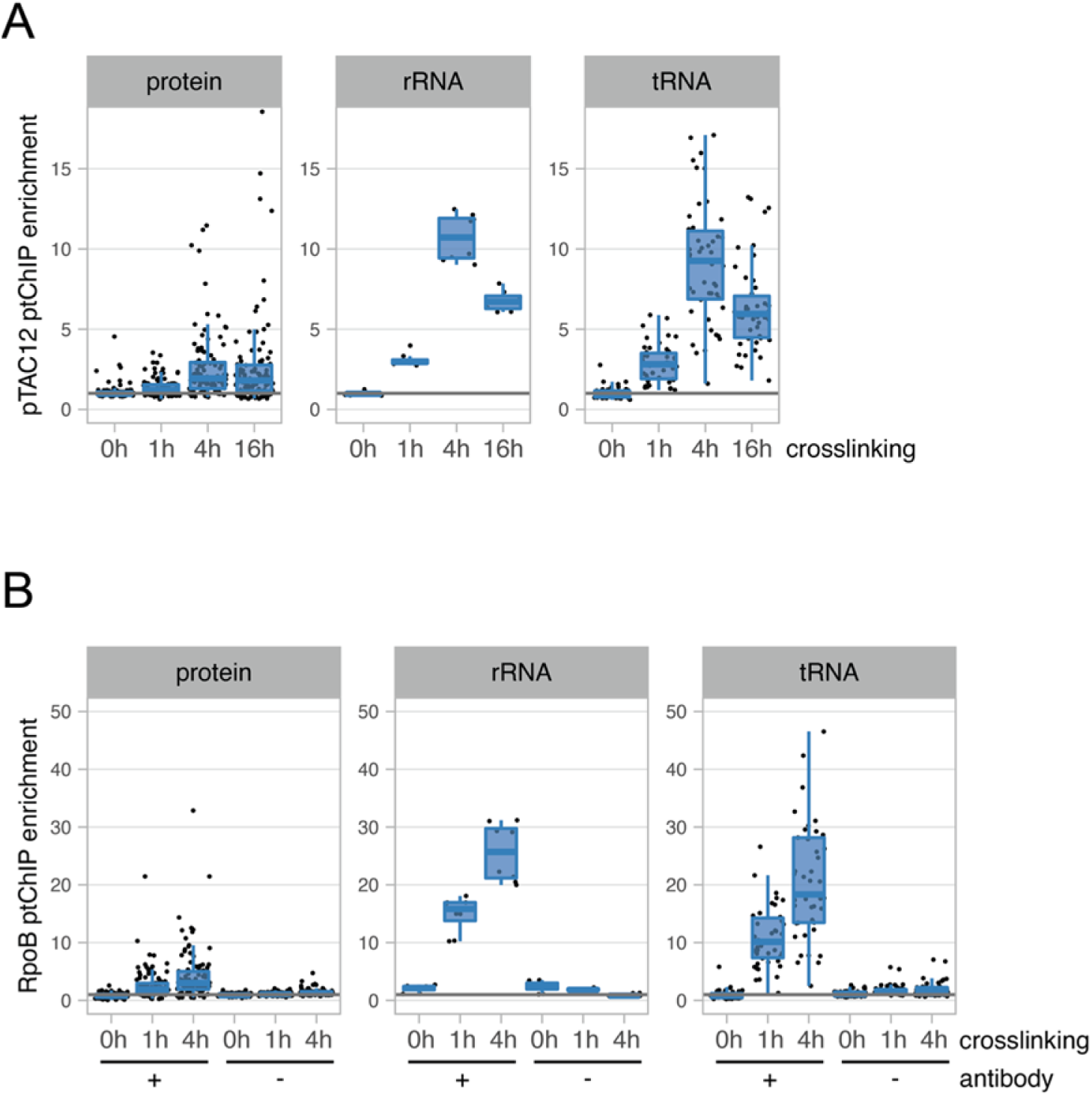
Detection of PEP binding to DNA using ptChIP-seq. A. Optimization of formaldehyde crosslinking time in ptChIP-seq. ptChIP-seq was performed using αHA antibody in plants expressing pTAC12-HA (Galvão et al., 2012) with no crosslinking or crosslinking of enriched chloroplasts with 4% formaldehyde for 1h, 4h and 16h. ptChIP-seq signals on annotated genes were calculated by dividing RPM normalized read counts from αHA ptChIP-seq in pTAC12-HA by RPM normalized read counts from αHA ptChIP-seq in Col-0 wild type. B. Optimization of formaldehyde crosslinking time and negative controls in ptChIP-seq. ptChIP-seq was performed using αRpoB antibody in Col-0 wild type plants with no crosslinking or crosslinking of enriched chloroplasts with 4% formaldehyde for 1h and 4h. ptChIP-seq was performed with and without the αRpoB antibody. ptChIP-seq signals on annotated genes were calculated by dividing RPM normalized read counts from αRpoB ptChIP-seq in Col-0 wild type by RPM normalized read counts from input samples. In A and B ptChIP-seq enrichments were calculated by dividing signal level on individual genes by the median signal level on genes in the *rpoB* operon, which is not transcribed by PEP and represents background signal levels. Genes were divided by the functions of their products into protein-coding, rRNA genes and tRNA genes. Average enrichments from two or three independent biological replicates are shown. Individual biological replicates are shown in Fig. S1.

### ptChIP-seq with αRpoB antibody is specific

Reliance of ptChIP-seq signal on formaldehyde crosslinking (Fig. 1AB) offers one line of evidence that this method is specific. Additionally, specificity of ptChIP-seq is supported by the lack of signal enrichment in controls without an antibody (Fig. 1B, Fig. S1B). To further test ptChIP specificity, we compared ptChIP-seq using αRpoB antibody in Col-0 wild-type to ptChIP-seq using αHA antibody in plants expressing pTAC12-HA (Galvão et al., 2012). Obtained ptChIP-seq enrichments were highly and significantly correlated between the two experiments when analyzed on annotated genes (Fig. 2A) or bins distributed throughout the entire plastid genome (Fig. S2). This indicates that pTAC12 and RpoB bind the same loci, as expected, and further supports high specificity of ptChIP-seq.

**Figure 2.**
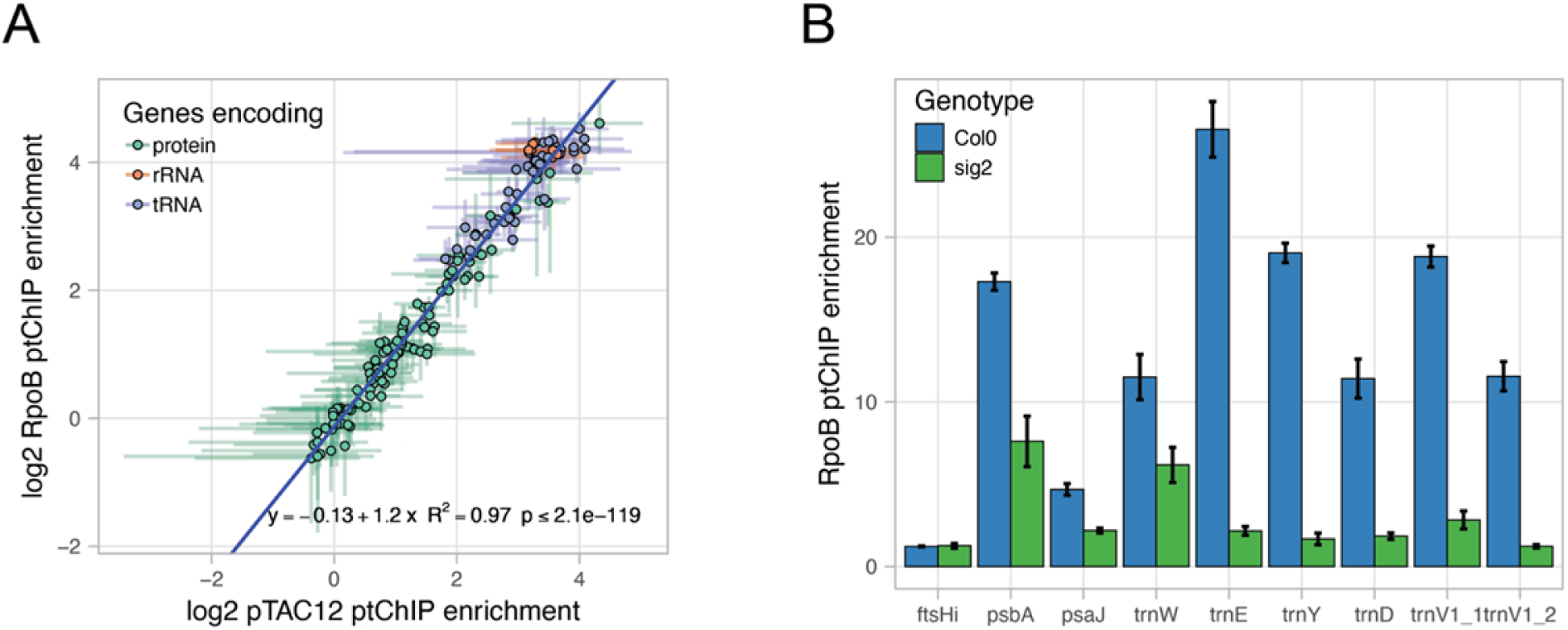
Specificity of ptChIP-seq with αRpoB antibody. A. pTAC12 and RpoB ptChIP-seq signals are highly correlated. Enrichment levels on annotated genes were compared between ptChIP-seq experiments using αHA antibody in plants expressing pTAC12-HA and using αRpoB antibody in Col-0 wild type plants. Data points are color-coded by function of the corresponding genes and show averages from three independent replicates. Error bars indicate standard deviations. Blue line represents the linear regression model. B. RpoB ptChIP-seq signal is reduced in *sig2* mutant on genes known to be affected by SIG2. Enrichment levels of ptChIP-seq using αRpoB antibody in Col-0 wild type and *sig2* were calculated on individual genes. Bars show averages from three independent biological replicates. Error bars indicate standard deviations.

A critical element of ChIP is a proper negative control. A genotype not expressing the epitope captures most sources of non-specific signal. For αHA ptChIP-seq in plants expressing pTAC12-HA, Col-0 wild-type serves as a proper negative control. Such a control is however much more difficult to obtain for αRpoB ptChIP-seq as the *rpoB* mutant is non-autotrophic (Allison et al., 1996). Because of that, αRpoB ptChIP-seq signal may instead be compared to input samples. The strong correlation between RpoB and pTAC12 ptChIP-seq experiments (Fig. 2A, Fig. S2A) indicates that input serves as a good negative control and that the αRpoB antibody may be used for ptChIP-seq.

To further test if αRpoB ptChIP-seq is specific, we investigated αRpoB ptChIP-seq signal on a NEP-transcribed negative control locus *ftsHi*/*ycf2* (Swiatecka-Hagenbruch et al., 2007). No enrichment was observed on *ftsHi* in Col-0 wild type (Fig. 2B), which indicates that ptChIP-seq is specific. Then we examined if PEP binding to DNA is affected in a mutant defective in SIG2, a sigma factor known to affect specific genes in Arabidopsis seedlings (Chi et al., 2015; Lerbs-Mache, 2011). We performed αRpoB ptChIP-seq in 4-day-old seedlings of Col-0 wild-type and *sig2* mutant. Because plastids are difficult to isolate from seedlings at this growth stage, we applied 4% formaldehyde for 4 h to the intact seedlings to capture protein-DNA interaction. RpoB enrichment on DNA in 4-day-old wild type seedlings was well correlated with that observed in 14-day-old seedlings (Fig. S2B), indicating that ptChIP can be applied to different developmental stages. We further analyzed the mean ptChIP-seq enrichment on a subset of loci that have previously been assayed for changes in RNA accumulation in *sig2* (Hanaoka et al., 2003; Kanamaru et al., 2001; Nagashima et al., 2004; Privat et al., 2003). ptChIP-seq enrichment was strongly reduced on *trnE*, *trnY*, *trnD*, and *trnV* (Fig. 2B), which is consistent with reported substantial reduction of steady state levels of those tRNAs in *sig2* (Hanaoka et al., 2003; Kanamaru et al., 2001; Nagashima et al., 2004; Privat et al., 2003). ptChIP-seq enrichment on *trnW* was reduced to 0.53 of Col-0, which is consistent with a small effect of *sig2* on the accumulation of its tRNA product (Kanamaru et al., 2001). ptChIP-seq enrichment on *psbA* was reduced to 0.44 of Col-0, consistently with a relatively small impact of *sig2* on the accumulation of its mRNA (Kanamaru et al., 2001). ptChIP-seq enrichment was also reduced on *psaJ*, which is consistent with prior RNA accumulation data (Nagashima et al., 2004). This indicates that αRpoB ptChIP results in *sig2* are generally consistent with prior RNA accumulation studies. Together, these results indicate that ptChIP-seq with αRpoB antibody is highly specific and may be used to assay the interactions of PEP with DNA.

### Complex pattern of PEP binding to DNA

Analysis of PEP binding across the plastid genome revealed a complex pattern of occupancy with preferential binding to genes encoding rRNA, tRNA, and some protein-coding genes (Fig. 3A). Most PEP binding is present within the inverted repeats (IR) where rRNA genes are located and in the large single copy region (LSC). The small single copy region (SSC) had little PEP binding (Fig. 3A). These observations are consistent with prior ChIP-chip study in tobacco (Finster et al., 2013) and assays on a limited number of loci in Arabidopsis (Yagi et al., 2012). Interestingly, we observed over a 20-fold dynamic range of ptChIP-seq signals between regions with various levels of PEP binding (Fig. 3A). This is also consistent with prior reports of PEP binding (Finster et al., 2013; Yagi et al., 2012) and transcription (Deng et al., 1987) and confirms that the levels of transcription may be greatly variable between individual genes.

**Figure 3.**
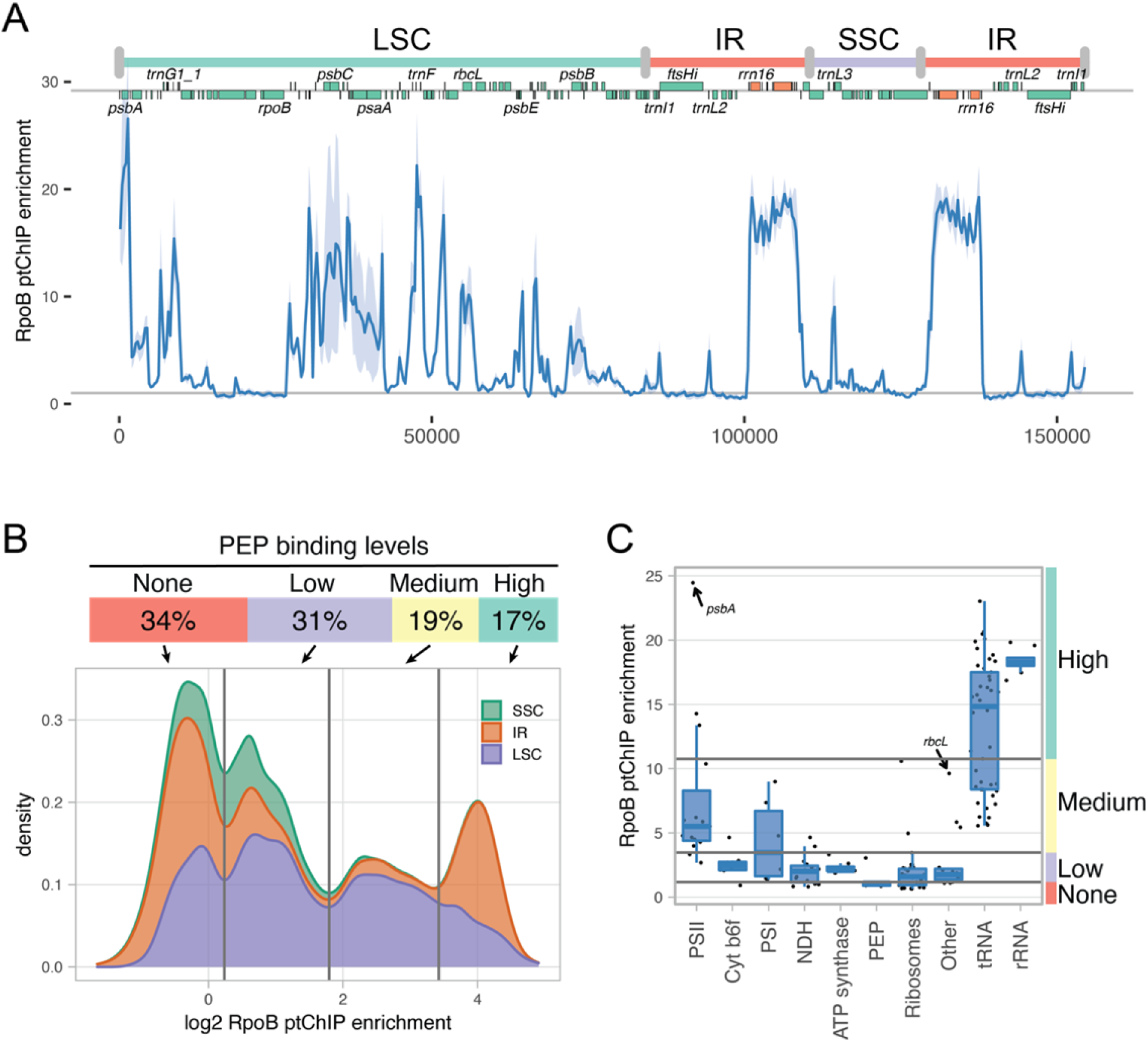
Complex pattern of PEP binding to plastid DNA. A. Genome-wide pattern of PEP binding to DNA. Signal enrichment from ptChIP-seq using αRpoB antibody in Col-0 wild type plants was calculated in 50 bp genomic bins and plotted throughout the entire plastid genome. Genome annotation including genomic regions, positions of annotated genes and names of selected individual genes are provided on top of the plot. Average enrichments from three independent biological replicates are shown. Light blue ribbon indicates standard deviation. B. Four preferred levels of PEP binding to DNA. Density plot of signal enrichments of ptChIP-seq using αRpoB antibody in Col-0 wild type plants. Average enrichments in 50 bp genomic bins from three independent biological replicates were analyzed in the SSC, IR and LSC. PEP binding level groups were determined by positions of local minima on the density plot. Percentages on top indicate fraction of all genomic bins assigned to a particular group. C. PEP binding to DNA of genes classified by the function of their products. Enrichment levels of ptChIP-seq using αRpoB antibody in Col-0 wild type were plotted on annotated genes split by the functions of gene products (Chotewutmontri and Barkan, 2018). PEP binding level groups are indicated on the right. Data points show averages from three independent biological replicates. Independent replicates are shown in Fig. S3.

We next tested if various levels of PEP binding are equally likely throughout the plastid genome. Distribution of ptChIP enrichment levels was multimodal with four peaks corresponding to no detectable PEP binding and three levels of PEP presence on the genome (Fig. 3B). Thirty four percent of the genome had no PEP binding, 31% had a low level of PEP, 19% had a medium level of PEP, and 17% had a high level of PEP (Fig. 3B). Focusing on annotated genes only, no PEP was detected on the *rpoB* operon, a subset of genes encoding ribosomal proteins, and a subset of genes encoding NDH subunits (Fig. 3C, Fig. S3A). Low and medium levels of PEP were detected on most genes encoding photosynthetic proteins and a subset of tRNA genes (Fig. 3C, Fig. S3A). High levels of PEP were found on the remaining tRNA genes, rRNA genes, and three genes encoding photosystem II subunits, including *psbA* (Fig. 3C, Fig. S3A). The multimodal distribution of PEP binding to DNA may be speculatively interpreted as an indication that the transcriptional machinery may adopt four distinct functional states corresponding to the four preferred levels of PEP binding.

Together, these results show a complex pattern of PEP binding to the plastid genome and suggest the presence of at least four preferred functional states of plastid transcriptional machinery.

### Dual impact of *sig2* and *sig6* mutants on PEP binding

Out of six sigma factors in Arabidopsis, only SIG2 and SIG6 are essential for proper chloroplast development (Chi et al., 2015; Lerbs-Mache, 2011). To determine the impact of those two sigma factors on PEP binding to DNA, we analyzed RpoB ptChIP-seq in *sig2* and *sig6* across the entire genome. The pattern of PEP binding was disrupted throughout the genome with most genes showing partial reduction in both mutants (Fig. 4A). The *sig6* mutant had a much stronger effect than *sig2* mutant on most PEP-bound loci (Fig. 4A)

**Figure 4.**
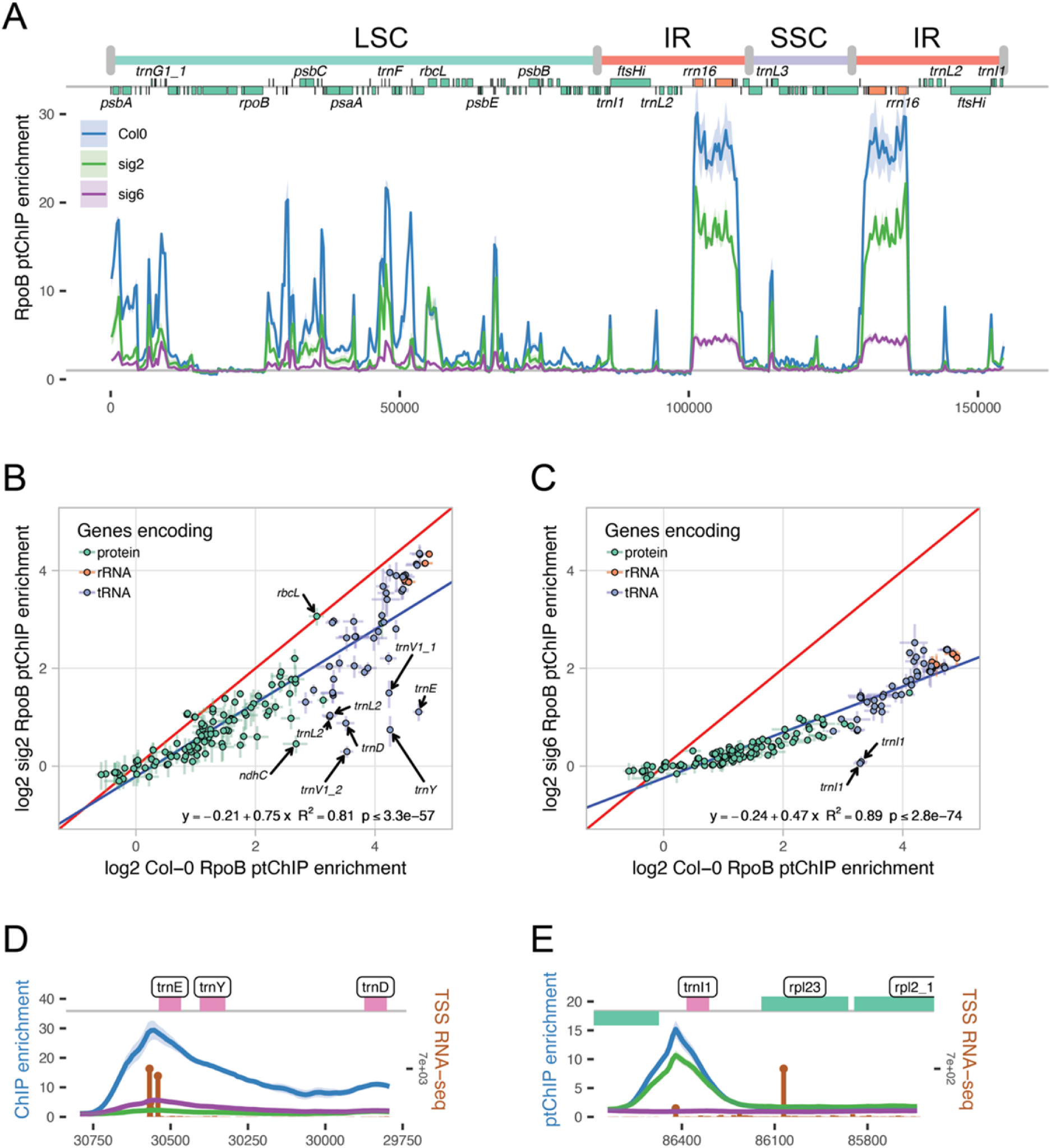
Dual impact of *sig2* and *sig6* mutants on PEP binding to plastid DNA. A. Genome-wide impact of *sig2* and *sig6* on of PEP binding to DNA. Signal enrichments from ptChIP-seq using αRpoB antibody in Col-0 wild type, *sig2* and *sig6* plants were calculated in 50 bp genomic bins and plotted throughout the entire plastid genome. Genome annotation including genomic regions, positions of annotated genes, and names of selected individual genes are provided on top of the plot. Average enrichments from three independent biological replicates are shown. Ribbons indicates standard deviations. B. Dual impact of *sig2* on PEP binding. Enrichment levels on annotated genes from ptChIP-seq using αRpoB antibody were compared between Col-0 wild type and *sig2* plants. C. Dual impact of *sig6* on PEP binding. Enrichment levels on annotated genes from ptChIP-seq using αRpoB antibody were compared between Col-0 wild type and *sig6* plants. In B and C data points are color-coded by the function of the corresponding genes and show averages from three independent replicates. Error bars indicate standard deviations. Blue line represents the linear regression model. Red line represents values equal between both genotypes. D. Reduction of PEP binding to DNA in *sig2* and *sig6* on the *trnEYD* operon. E. Reduction of PEP binding to DNA in *sig2* and *sig6* on *trnI1*. In D and E signal enrichments from ptChIP-seq using αRpoB antibody in Col-0 wild type, *sig2* and *sig6* plants were calculated in 10 bp genomic bins and plotted at the relevant locus. Color coding of ptChIP-seq data corresponds to data shown in Fig. 4A. Average enrichments from three independent biological replicates are shown. Ribbons indicates standard deviations. Brown vertical lines indicate sense strand data from three combined replicates of TSS RNA-seq. Genome annotation is shown on top.

Regression analysis of *sig2* compared to Col-0 wild type revealed that ptChIP-seq signals in Col-0 wild type and *sig2* are significantly correlated with a slope of 0.75 (Fig. 4B). This indicates that in *sig2*, most genes have a consistent, moderate reduction of PEP binding (Fig. 4B). Only a few genes had PEP binding reduced to much greater extents than indicated by the genome-wide trend. These included four previously studied tRNA genes (Fig. 2B, Fig. 4AD), *trnL2* (Fig. 4B, Fig. S4A), and one protein-coding gene *ndhC* (Fig. 4B, Fig. S4A). However, PEP binding was unchanged on *rbcL* (Fig. 4B, Fig. S4A).

A similar pattern was observed in *sig6* where PEP binding was strongly reduced throughout the genome. PEP binding in *sig6* and Col-0 wild type remained significantly correlated, but the slope of the regression line was reduced to 0.47 (Fig. 4C). Only one gene had an almost complete loss of PEP binding, the tRNA gene, *trnI*, located in the inverted repeat (Fig. 4CE). Consistently, accumulation of the *trnI* tRNA product was significantly reduced in *sig6* (Fig. S4B).

Decreased PEP binding in *sig2* and *sig6* mutants may be due to lower abundance of PEP. To test this possibility, we assessed the protein accumulation of a PEP subunit, RpoC1 (Fig. S4C). The accumulation levels of this subunit were slightly lower in both *sig2* and *sig6* compared to the Col-0 wild type (Fig. S4C), which indicates a small reduction of PEP abundance. These results suggest that SIG2 and SIG6 have specific impacts on limited numbers of genes together with weaker but broad impacts on PEP occupancy throughout the genome.

### PEP preferentially binds to gene promoters

Previous studies identified preferential binding of PEP to two promoters of photosystem II genes (Ding et al., 2019; Yagi et al., 2012). To determine if promoter binding is a more general property of PEP, we performed RNA-seq designed to identify triphosphorylated 5’ ends of primary transcripts (Zhelyazkova et al., 2012), or TSS RNA-seq, and compared it with the ptChIP-seq signal on PEP-transcribed genes with known promoter locations. Consistent with prior findings (Ding et al., 2019; Yagi et al., 2012), PEP binding was strongly enriched on annotated *psbA* and *psbEFLJ* promoters (Fig. 5A). These promoters also had strong TSS RNA-seq signals (Fig. 5A), which confirms that peaks of PEP binding coincide with transcription start sites. We observed similar preferential PEP binding to other known promoters including *psaA* and *rbcL* (Fig. 5B, Fig. S5A). When averaged over all promoters previously identified in Arabidopsis (Fig. S5AB), PEP binding and TSS RNA-seq signals were strongly enriched on gene promoters (Fig. 5C). These results indicate that preferential binding to gene promoters may be a general property of PEP.

**Figure 5.**
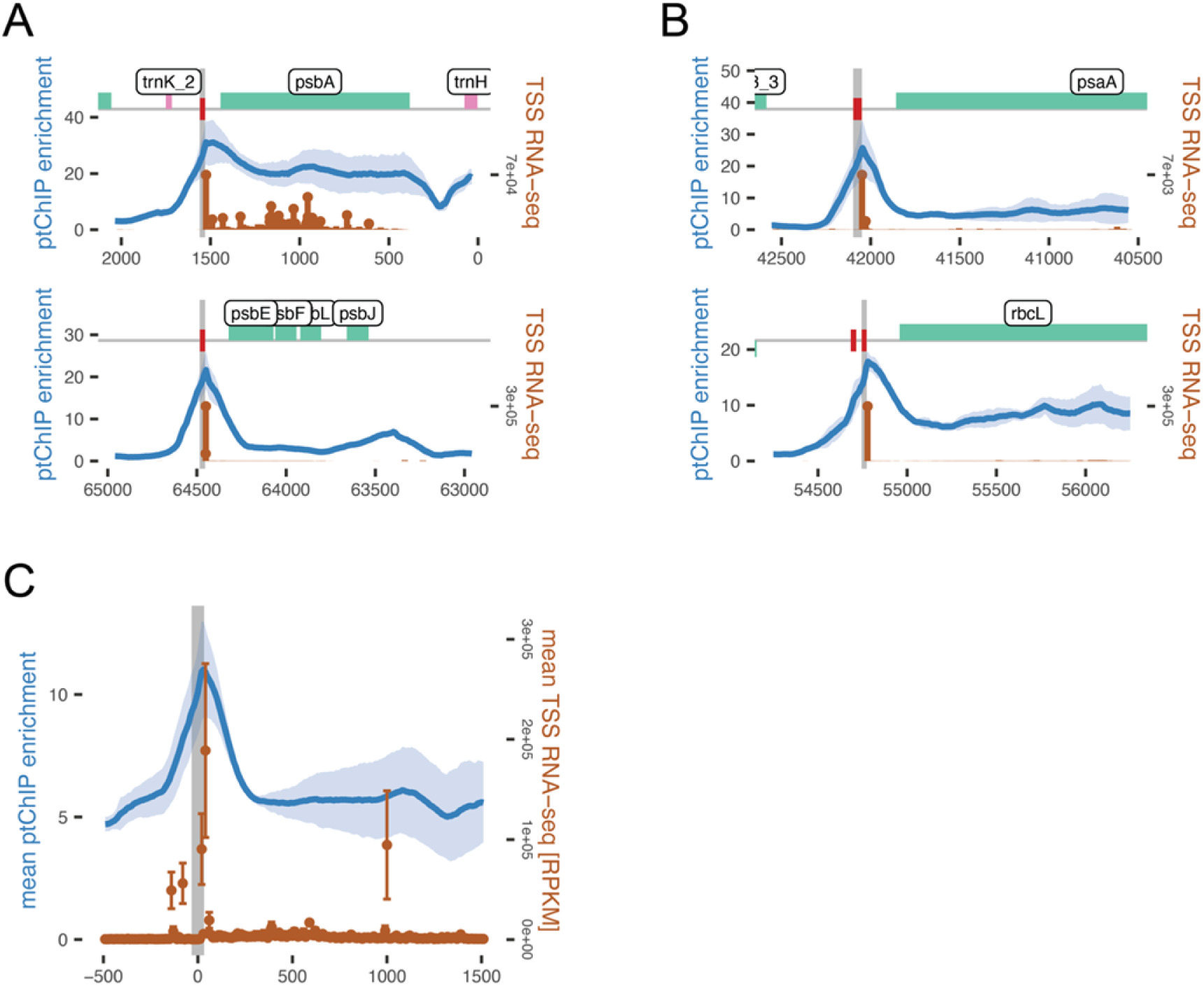
Preferential binding of PEP to gene promoters. A. Binding of PEP and transcription start sites on promoters previously shown to be bound by PEP. B. Preferential binding of PEP and transcription start sites on other promoters. In A and B signal enrichment from ptChIP-seq using αRpoB antibody in Col-0 wild type was calculated in 10 bp genomic bins and plotted at the relevant loci. Average enrichments from three independent biological replicates are shown. Light blue ribbons indicate standard deviations. Brown vertical lines indicate sense strand data from three combined replicates of TSS RNA-seq. Grey vertical line indicates position of the annotated promoter. Genome annotation is shown on top. C. Average binding of PEP and transcription start sites on all promoters found in Arabidopsis. Genomic regions surrounding all PEP promoters we identified in prior studies in Arabidopsis were aligned and mean ptChIP-seq enrichment using αRpoB antibody in Col-0 wild type was calculated in 10 nt genomic bins for each biological replicate. Mean value from three biological replicates is shown. Light blue ribbon corresponds to standard deviation. Mean TSS RNA-seq signal was calculated from both strands in 10 bp genomic bins for each biological replicate. Brown dots correspond to mean values from three biological replicates. Error bars indicate standard deviations. Grey vertical line indicates aligned position of the annotated promoter.

### PEP binding is correlated with steady state levels of RNA

Posttranscriptional regulation has a major impact on plastid gene expression (Barkan, 2011). Therefore, the levels of PEP binding to DNA may have a limited correlation with steady state levels of RNA. To test this prediction, we split the genome into 250nt bins and counted average TSS RNA-seq read counts and ptChIP-seq enrichments from three replicates of both experiments. Inverted Repeat regions were not included because highly structured regions within mature rRNAs may inhibit Terminator exonuclease. PEP binding to DNA and steady state levels of primary transcripts were significantly correlated (Fig. 6A). This suggests that differences in steady state levels of RNA may to some extent be explained by differences in PEP transcription.

**Figure 6.**
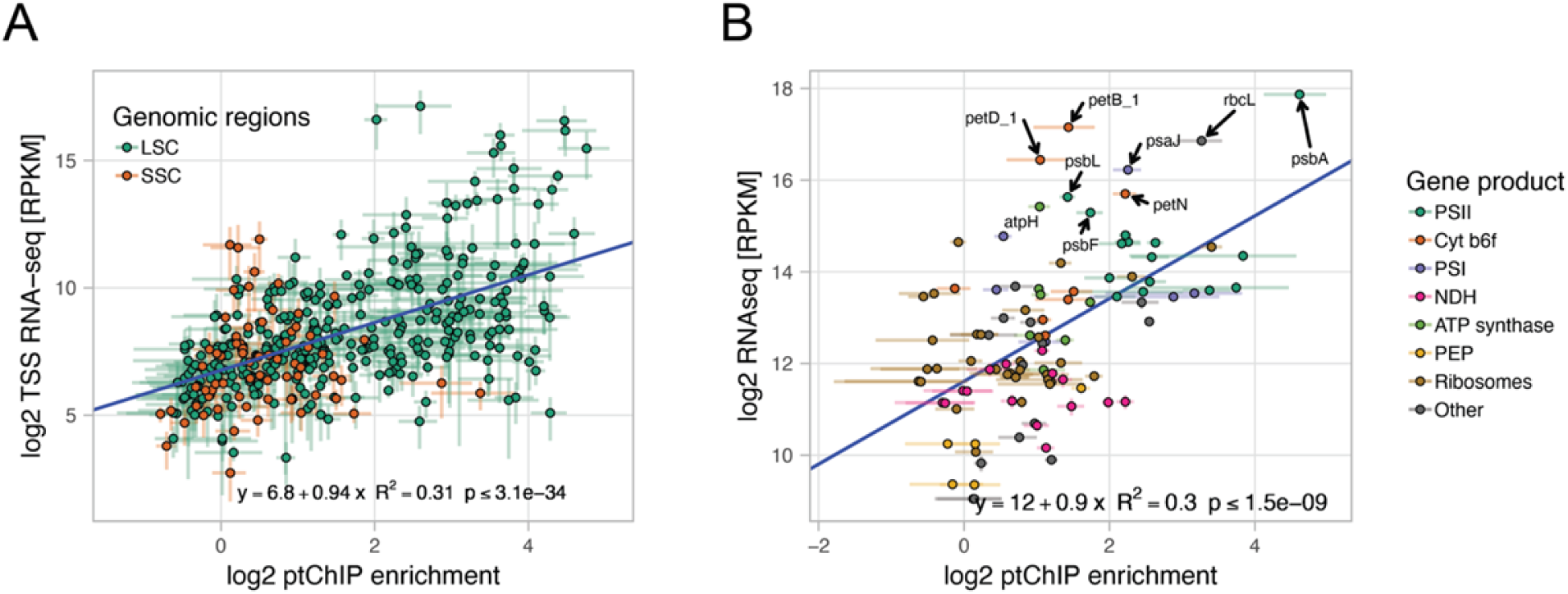
Correlation of PEP binding with steady state levels of RNA. A. RpoB ptChIP-seq enrichments and TSS RNA-seq signals are highly correlated. Enrichment levels of RpoB ptChIP-seq and RPKM normalized TSS RNA-seq signals in Col-0 wild type were compared on 250 bp genomic bins including LSC and SSC. Data points are color-coded by locations. Blue line represents the linear regression model. B. RpoB ptChIP-seq enrichments and total RNA-seq signals are highly correlated on protein coding genes. Enrichment levels of RpoB ptChIP-seq in 14 days old Col-0 wild type plants (this study) and RPKM normalized total RNA-seq signals from a similar developmental stage (Thieffry et al., 2020) were compared on annotated protein coding genes. Data points are color-coded by the function of the corresponding genes and show averages from three independent replicates. Error bars indicate standard deviations. Blue line represents the linear regression model.

Transcription start sites are located outside of annotated genes. Consistently, there are many genomic bins which have high levels of PEP binding to DNA and low levels of TSS RNA-seq (Fig. 6A). To overcome this limitation of TSS RNA-seq, we used a previously published RNA-seq dataset from 14-day-old plants (Thieffry et al., 2020) and compared the steady state level of total RNA to the level of PEP binding to DNA on annotated genes. RNA-seq levels on rRNA and tRNA genes were not correlated with PEP binding (Fig. S6A), probably due to ribodepletion of RNA samples prior to the library prep and highly structured tRNAs being poor substrates for library production. However, annotated protein-coding genes had a significant correlation between RNA-seq and ptChIP-seq signals (Fig. 6B). R^2^ of 0.3 indicates that about a third of RNA-seq variance may be predicted by the level of PEP binding to DNA. The reminder of RNA-seq variance is likely affected by RNA processing and degradation, and a subset of PEP-transcribed mRNAs, such as *psbA*, *rbcL*, and *petB*, appears to be particularly stable (Fig. 6B). Together, these results indicate that PEP binding to DNA, which may be interpreted as a proxy for the level of transcription, has a significant impact on the steady state levels of RNA and possibly, more generally, on gene expression.

## DISCUSSION

### Detection of protein-DNA interactions in plastids

Several lines of evidence support the specificity of ptChIP-seq in detecting PEP binding to plastid DNA. A key feature of this protocol is efficient formaldehyde crosslinking of enriched chloroplasts combined with stringent immunoprecipitation conditions. Together, they allow detection of only protein-DNA interactions that have been preserved by crosslinking. This is in stark difference to some other protein-nucleic acid interaction studies in plastids (Barkan, 2009; Newell and Gray, 2010). While a long formaldehyde treatment may increase the risk of crosslinking artifacts (Walker et al., 2020), our data clearly demonstrate no RpoB ptChIP-seq signal on loci that are not transcribed by PEP, such as the *rpoB* operon and the coding region of *ftsHi*/*ycf2* (Hajdukiewicz et al., 1997; Swiatecka-Hagenbruch et al., 2007). This indicates that signal preserved by crosslinking is very likely to be specific.

It is also important to note that our protocol can capture the pattern of PEP-DNA interactions in both enriched mature chloroplasts and intact seedlings with similar specificity. Transcriptional regulation is particularly important in the early stages of chloroplast differentiation (Pfannschmidt et al., 2015). Considering the difficulty of plastid isolation at early growth stages, our ptChIP-seq protocol for intact seedlings can be a powerful approach to explore the mechanism of transcriptional regulation in developing chloroplasts.

ptChIP-seq with the αRpoB antibody shows a high dynamic range of PEP binding to DNA, which is consistent with prior observations on small subsets of specific loci in Arabidopsis and tobacco (Finster et al., 2013; Yagi et al., 2012). The dynamic range of ptChIP-seq is however an order of magnitude higher than previously reported ChIP-chip study of RpoA in tobacco (Finster et al., 2013). This indicates that ptChIP-seq may substantially expand our understanding of PEP transcription and overall regulation of plastid gene expression.

### Complexity of PEP transcription

The pattern of PEP binding to DNA is complex, which illustrates that variable levels of protein production are at least partially caused by variable levels of PEP transcription. Although the involvement of NEP transcription remains unknown, the complex pattern of PEP binding is not consistent with a simplistic view of the model assuming full transcription of the plastid genome (Shi et al., 2016).

Intensity of PEP binding to DNA throughout the plastid genome is not distributed normally and instead shows four preferred levels. We speculate that these levels correspond to four preferred functional states of PEP. These four states could be caused by specific cis-acting elements and regulatory proteins bound to those elements. They could also be reflected by preferred interactions between the core PEP complex and accessory proteins, or even could be an indication that the genome exists in four favored structural states, which would be reminiscent of the situation in the nucleus (Roudier et al., 2011).

A strong preferential binding of PEP within gene promoters, around transcription start sites, is consistent with the concept of RNA polymerase pausing on gene promoters (Landick, 2006). While it has been previously shown on the *psbEFLJ* operon (Ding et al., 2019), our results indicate that pausing may be a general property of PEP. The role of PEP pausing in transcription or gene regulation remains unresolved.

### Role of SIG2 and SIG6 in PEP recruitment

PEP is recruited to its promoters by binding of sigma factors in a sequence specific manner. Among six nuclear-encoded sigma factors in Arabidopsis, SIG1, SIG3, SIG4 and SIG5 have highly specific functions and are not required for proper chloroplast development. In contrast, SIG2 and SIG6 have more general impacts on plastid transcription and are required for early chloroplast development (Börner et al., 2015; Chi et al., 2015; Puthiyaveetil et al., 2021).

Strong impacts of *sig2* and *sig6* on a limited number of tRNA genes accompanied by a genome-wide moderate reduction of PEP binding could be explained by complex patterns of SIG2 and SIG6 binding specificities. In this scenario, strong non-redundant impacts on a few targets would be accompanied by weaker non-redundant but still specific roles on all remaining PEP-transcribed genes. We, however, propose a simpler explanation where SIG2 and SIG6 have a dual impact on PEP binding by a combination of direct and indirect mechanisms. In this model, both SIG2 and SIG6 directly and non-redundantly only impact limited numbers of mostly tRNA genes. Then, tRNA deficiencies negatively impact plastid translation and to some extent, PEP production, which leads to consistent reductions of PEP occupancy throughout the genome. This model is consistent with the observation that *sig2* showed a stronger decrease of tRNA compared to mRNA for photosynthesis-associated genes (Kanamaru et al., 2001) and also explains why *sig2* and *sig6* mutants recover from early developmental defects and produce fully functional chloroplasts later in development (Ishizaki et al., 2005; Privat et al., 2003).

All tRNAs strongly affected in *sig2* or *sig6* mutants lose PEP binding not only throughout their transcribed sequences but also on their promoters. This result suggests that pausing is tightly coupled with transcription elongation in plastids.

### Contribution of transcription to gene regulation in plastids

Significant correlation between PEP binding to DNA determined by ptChIP-seq and steady state levels of RNA determined by RNA-seq allows estimation of the contribution of PEP binding to RNA accumulation. We estimate that about 30% of RNA accumulation can be explained by PEP binding. PEP binding may be interpreted as a proxy for transcription rates, although it should be noted that PEP elongation rates remain unknown and may be variable between various genes. Several prior run on and ChIP studies also suggest that transcription has a significant role in gene regulation (Deng et al., 1987; Finster et al., 2013). It is also supported by the observation that reduction of PEP recruitment in sigma factor mutants is comparable to previously reported reductions of RNA levels. However, this is to our knowledge the first study providing a genome-wide quantitative estimate of the relationship between transcription and RNA levels.

Our observations about the contribution of transcription to gene regulation apply to the variability between genes. Changes in transcription between developmental or environmental conditions have mixed impacts on gene expression (Isono et al., 1997; Krupinska and Apel, 1989; Shiina et al., 1998). The presented approach may be used in future studies to uncover the contribution of PEP to condition-dependent plastid gene regulation.

## METHODS

### Plant materials and growth conditions

*Arabidopsis thaliana* wild-type Columbia-0 (Col-0) ecotype was used in all analyses. We used the following genotypes: *sig2-2* (SALK_045706) (Woodson et al., 2012), *sig6-1* (SAIL_893_C09) (Ishizaki et al., 2005) and pTAC12-HA (*HMR::HA*/*hmr-5*) transgenic line (Galvão et al., 2012). For experiments with 14-day-old plants, seeds were stratified in darkness at 4°C for 48 hours and grown on soil at 22°C under white LED light (100 μmol m^-2^ s^-1^) in 16h/8h day/night cycle. For experiments with 4-day-old plants, seeds were stratified in darkness at 4°C for 48 hours and grown on 0.5 X MS plates (0.215% MS salts, 0.05% MES-KOH pH 5.7, 0.65% Agar) for four days at 22°C under constant white LED light (50 μmol m^-2^ s^-1^).

### Chloroplast enrichment and crosslinking

Chloroplasts from 14-day-old seedlings were enriched following the protocol described by Nakatani and Barber with minor modifications (Nakatani and Barber, 1977). In brief, 5 grams of rosette leaves were harvested and rinsed 3 times with ultra-pure water to eliminate soil debris and homogenized in chloroplast enrichment buffer (0.33 M Sorbitol, 30 mM HEPES-KOH (pH 7.5), 0.001% β-mercaptoethanol) using a blender. The homogenate was filtered through two layers of Miracloth, and the flow-through was centrifuged at 1500 g for 5 minutes at 4°C. The pelleted chloroplasts were resuspended in 1 ml of chloroplast enrichment buffer. Chlorophyll concentration was determined by resuspending 10 μl of the chloroplast fraction in 1 ml 80% acetone and measuring its absorbance at 652 nm as reported (Inskeep and Bloom, 1985). To cross-link DNA to proteins, 4% final concentration of formaldehyde was applied to the amount of chloroplast corresponding to 200 μg of chlorophyll followed by incubation at 4°C for 4h unless indicated otherwise. Formaldehyde was quenched by diluting the chloroplasts 5 times in the chloroplast enrichment buffer containing 125 mM glycine, followed by chloroplast pelleting at 1500 g at 4°C. For experiments using 4-day-old seedlings, whole seedlings were vacuum infiltrated with 4% formaldehyde for 10 min as reported previously (Rowley et al., 2013) and incubated for 4h at 4°C.

### ptChIP-seq

ptChIP-seq protocol was based on a previously published nuclear ChIP protocol (Rowley et al., 2013). In brief, enriched chloroplasts from 14-day-old plants corresponding to 50 μg chlorophyll were subject to *in vitro* crosslinking. Alternatively, 50 4-day-old seedlings were crosslinked *in vivo*, flash-frozen, homogenized in lysis buffer (50 mM Tris-HCl pH 8.0, 10 mM EDTA, 1% SDS) and filtered through two layers of Miracloth. Obtained samples were sonicated to achieve DNA fragments ranging from 200 nt to 300 nt using a QSonica Q700 sonicator. The fragmented samples were incubated overnight with 1 μg of monoclonal anti-HA antibody (Invitrogen catalog number 26183) or 5 μg of polyclonal anti-RpoB antibody (PhytoAB catalog number Phy1239) with 40 μl Protein G Dynabeads (Invitrogen catalog number 10004D) or 60 μl Protein A Dynabeads (Invitrogen catalog number 10002D) respectively. After incubation, the beads were washed, and DNA was eluted, and reverse cross-linked as described (Rowley et al., 2013). High throughput sequencing libraries were prepared as reported (Bowman et al., 2013) and sequenced using an Illumina NovaSeq 6000 S4 flow-cell with 150×150 paired-end configuration at the University of Michigan Advanced Genomics Core.

### TSS RNA-seq

Five micrograms of RNA isolated from enriched chloroplasts was digested with 6U of Terminator exonuclease (Lucigen catalog number TER51020) for 1h at 30°C in 20µl reaction volume. The reaction was stopped by addition of 1µl of 100mM EDTA. Subsequently, digested RNA was purified with acidic buffer-saturated phenol, washed with 70% ethanol, and resuspended in 10µl of water. Purified RNA (∼10ng) was submitted to the University of Michigan Biomedical Research Core where libraries were generated with SMARTer Stranded Total RNA-Seq Kit-Pico Input Mammalian and sequenced. Modifications to the original protocol include: no RNA fragmentation, no rRNA depletion and inclusion of size selection (∼200bp fragments) of the libraries before sequencing.

### Data analysis

The obtained raw sequencing reads were trimmed using trim_galore v.0.4.1, and mapped to the TAIR10 Arabidopsis plastid genome (www.arabidopsis.org) using Bowtie2 v.2.2.8 (Langmead and Salzberg, 2012). For ptChIP-seq reads were PCR de-duplicated using PICARD tools. Read counts on defined genomic regions were determined using bedtools v.2.25.0 (Quinlan and Hall, 2010). ptChIP-seq signals on annotated genes were calculated by dividing RPM normalized read counts from αHA or αRpoB ptChIP-seq by RPM normalized read counts from αHA ptChIP-seq in Col-0 wild type or input samples, respectively. ptChIP-seq enrichments on annotated genes were calculated by dividing signal levels on individual genes by the median signal level on genes in the *rpoB* operon, which is not transcribed by PEP and represents background signal levels. ptChIP-seq enrichments on genomic bins were calculated by dividing signal levels on individual bins by the signal level on the entire *rpoB* operon.

### Identification of promoters

PEP promoters in the plastid genome were identified based on previous studies representing promoter sequences in Arabidopsis (Favory et al., 2005; Fey et al., 2005; Hanaoka et al., 2003; Hoffer and Christopher, 1997; Ishizaki et al., 2005; Kanamaru et al., 2001; Liere et al., 1995; Nagashima et al., 2004; Privat et al., 2003; Shimmura et al., 2008; Sriraman et al., 1998; Swiatecka-Hagenbruch et al., 2007; Zghidi et al., 2007). Regions between -35 (TTGACA) and - 10 (TATAAT) consensus motives were identified and are shown in Figure S5.

### Immunoblot analysis

Total proteins were extracted from 4-day-old seedlings by incubating homogenized samples in the sample buffer (20 mM Tris-HCl (pH 6.8), 3% β-mercaptoethanol, 2.5% sodium dodecyl sulfate, 10% sucrose) with cOmplete protease inhibitor cocktail (Roche) for 1.5 h at room temperature. After removing debris by centrifugation, 20 µg proteins were separated by SDS-PAGE. To detect RpoC1, monoclonal anti-RpoC1 antibody (PhytoAB catalog number PHY1904) and anti-mouse IgG antibody conjugated with horseradish peroxidase (Amersham catalog number NA931) were used as the primary and secondary antibodies, respectively. Protein bands were visualized using chemiluminescence reagents (ECL Prime Western Blotting Detection Reagent, Amersham) and an ImageQuant LAS 4000 imager.

### tRNA quantification

Total RNA was isolated from 50 fresh 4 days-old seedlings from wild type Col-0 and *sig6-1* mutant plants using Trizol following the manufacturer protocol. Following DNaseI digestion, 500 and 1000 ng of RNA were used to generate primer specific (for tRNA-Ile-CAU) and polyA cDNA, respectively, using the SuperScript III First Strand Kit following the instructions for highly secondary structured templates. Real-time PCR was performed using the KAPA Sybr Green 2x kit with the following primers: Ath_Actin2_FWD

5’GAGAGATTCAGATGCCCAGAAGTC3’, Ath_Actin2_REV
5’TGGATTCCAGCAGCTTCCA3’, tRNA-Ile-CAU_FWD
5’ATCCATGGCTGAATGGTTAAAGCG3’, tRNA-Ile-CAU-REV
5’CATCCAGTAGGAATTGAACCTACGA3’. Results were analyzed using the ddCT method.

### Accession numbers

The sequencing data from this study have been submitted to the NCBI Gene Expression Omnibus (GEO; http://www.ncbi.nlm.nih.gov/geo/) under accession number GSE192568. Sequencing data presented in this study are available through a dedicated publicly available Plastid Genome Visualization Tool (Plavisto) at http://plavisto.mcdb.lsa.umich.edu.

## Acknowledgements

This work was supported by a grant from the National Science Foundation (MCB 1934703) to A.T.W. S.F. was supported by grants from the Japanese Society for the Promotion of Science (19J01779, 20K15819). The pTAC12-HA (*HMR::HA*/*hmr-5*) transgenic line was kindly provided by Meng Chen (University of California, Riverside).

## Author contributions

V.M.P., S.J, S.F. and J.K. performed research. V.M.P, S.F. and A.T.W analyzed data. A.T.W. wrote the paper.

## SUPPLEMENTAL FIGURES

**Figure S1.**
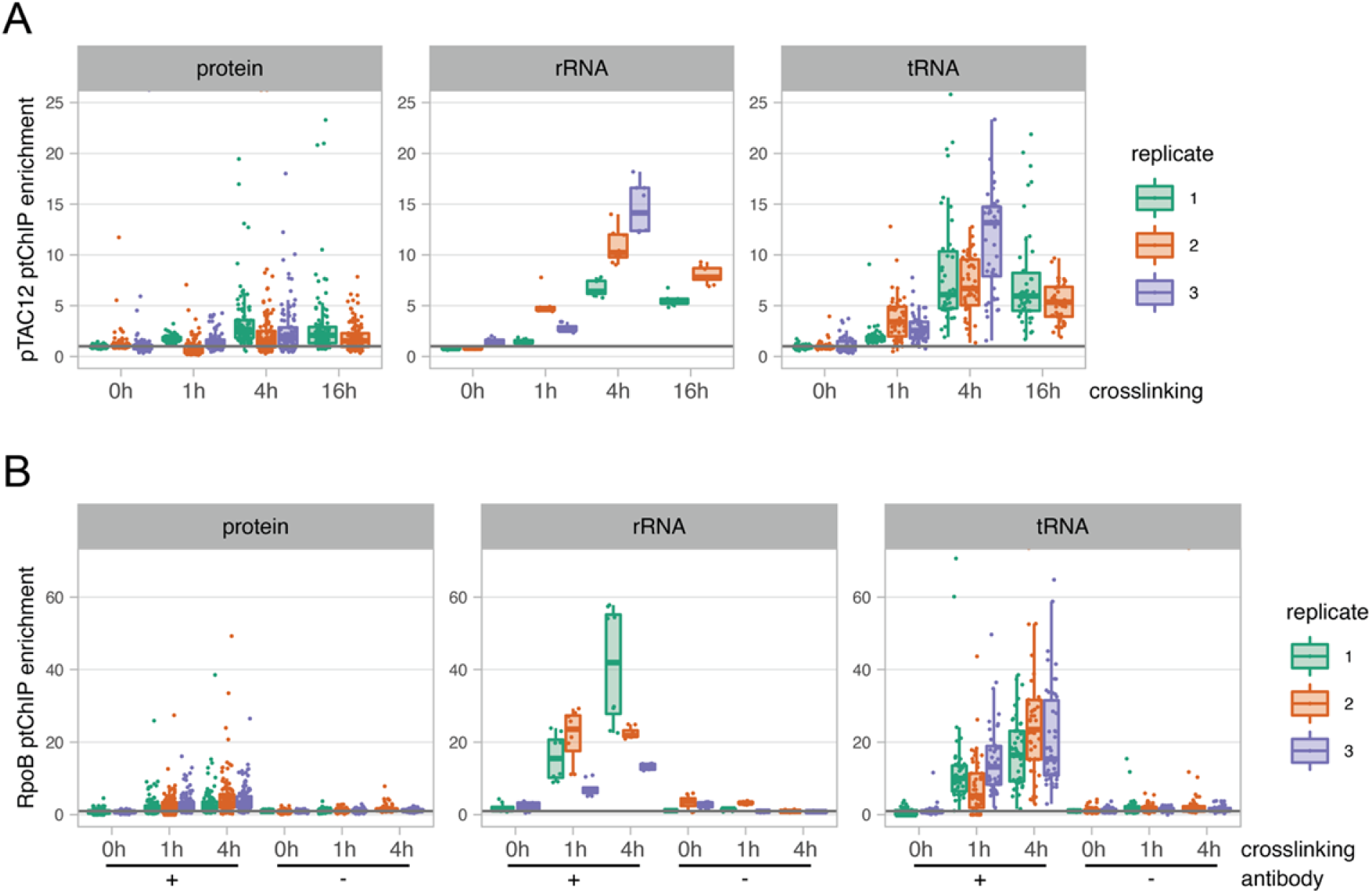
Detection of PEP binding to DNA using ptChIP-seq. Individual biological replicates of data shown in Fig. 1. A. Optimization of formaldehyde crosslinking time in ptChIP-seq. Individual biological replicates of ptChIP-seq performed using αHA antibody in plants expressing pTAC12-HA shown in Fig. 1A. B. Optimization of formaldehyde crosslinking time and negative controls in ptChIP-seq. Individual biological replicates of ptChIP-seq using αRpoB antibody in Col-0 wild type plants shown in Fig. 1B.

**Figure S2.**
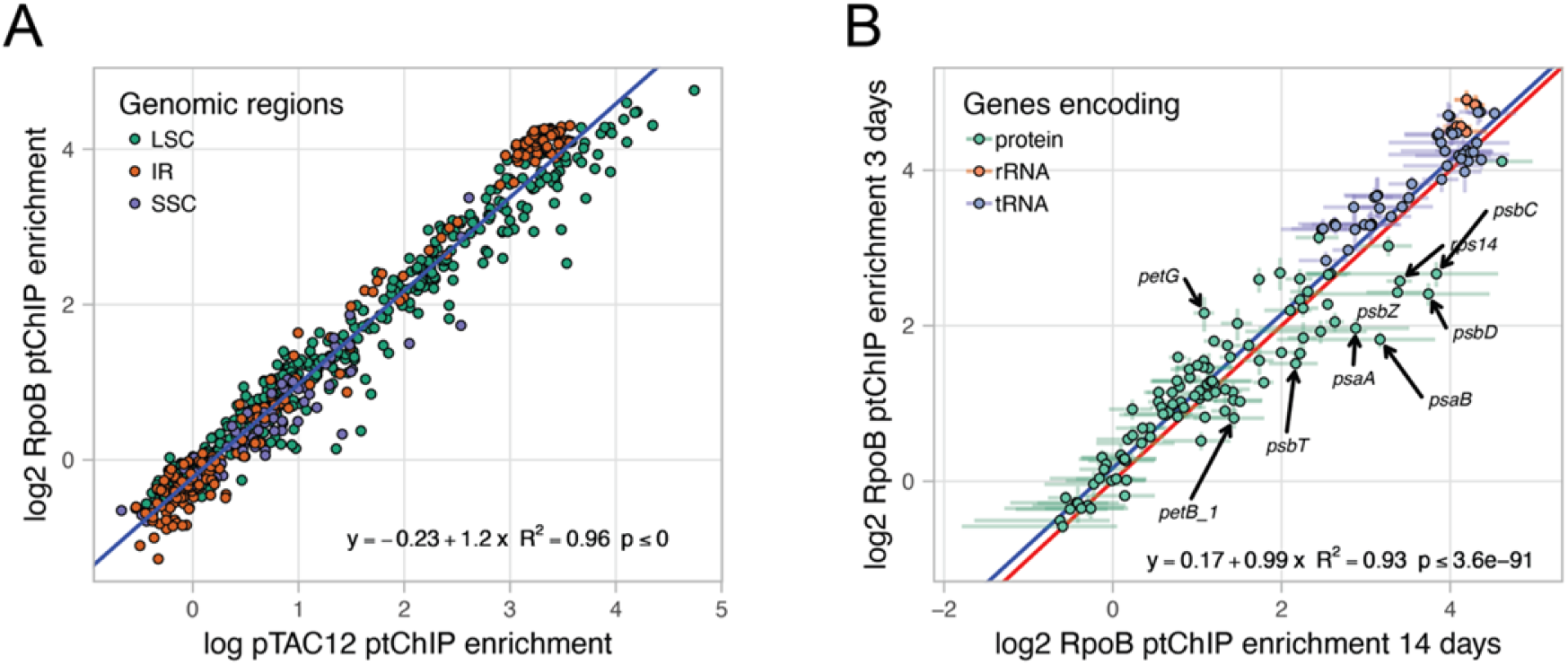
Specificity of ptChIP-seq with αRpoB antibody. A. pTAC12 and RpoB ptChIP-seq signals are highly correlated. Enrichment levels on 250 bp genomic bins were compared between ptChIP-seq experiments using αHA antibody in plants expressing pTAC12-HA and using αRpoB antibody in Col-0 wild type plants. Data points are color-coded by locations of bins within the LSC, IR or SSC. Blue line represents the linear regression model. B. RpoB ptChIP-seq signals are highly correlated between 4-day-old plants and 14-day-old plants. Enrichment levels on annotated genes from ptChIP-seq using αRpoB antibody in Col-0 wild type plants were compared between 4-day-old and 14-day-old plants. Data points are color-coded by the function of the corresponding genes and show averages from three independent replicates. Error bars indicate standard deviations. Blue line represents the linear regression model.

**Figure S3.**
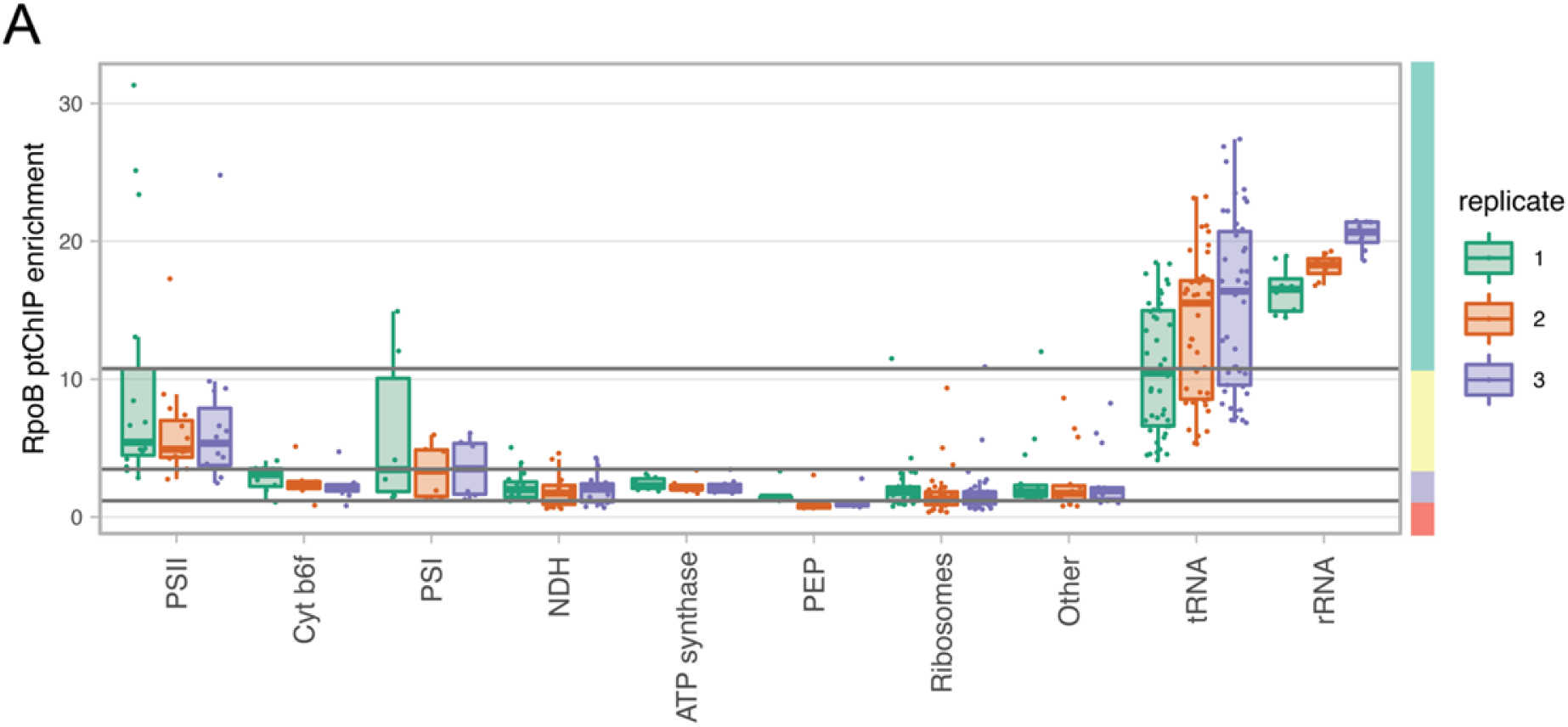
Complex pattern of PEP binding to plastid DNA. Individual biological replicates of data shown in Fig. 3C. A. PEP binding to DNA of genes classified by the function of their products. Enrichment levels of ptChIP-seq using αRpoB antibody in Col-0 wild type were plotted on annotated genes split by the function of gene products (Chotewutmontri and Barkan, 2018). PEP binding level groups are indicated on the right. Colors indicate data from independent replicates.

**Figure S4.**
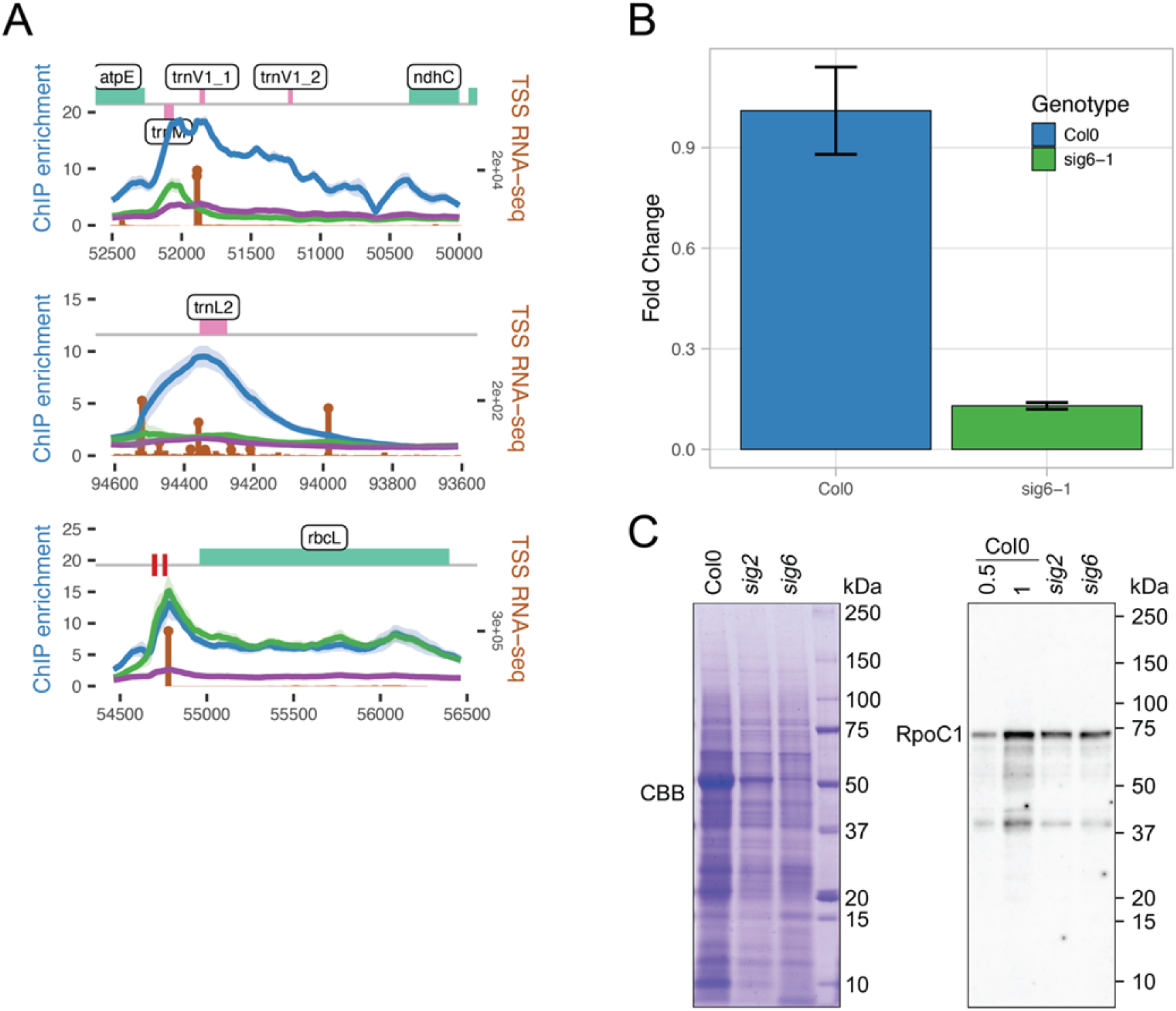
Dual impact of *sig2* and *sig6* mutants on PEP binding to plastid DNA. A. Reduction of PEP binding to DNA in *sig2* and *sig6* on *trnV*, *trnL* and *rbcL*. Signal enrichments from ptChIP-seq using αRpoB antibody in Col-0 wild type, *sig2* and *sig6* plants were calculated in 10 bp genomic bins and plotted at the relevant locus. Color-coding of ptChIP-seq data corresponds to data shown in Fig. 4A. Average enrichments from three independent biological replicates are shown. Ribbons indicates standard deviations. Brown vertical lines indicate sense strand data from three combined replicates of TSS RNA-seq. Genome annotation is shown on top. B. Reduction of the tRNA product of *trnI1* (tRNA-Ile-CAU) in the *sig6* mutant. Accumulation of the tRNA was assayed using RT-qPCR. Average and standard deviation from three biological replicates are shown. C. Abundance of the RpoC1 protein in *sig2* and *sig6*. 20 µg of total proteins were loaded. Gel stained with Coomassie brilliant blue (CBB) is shown as a loading control.

**Figure S5.**
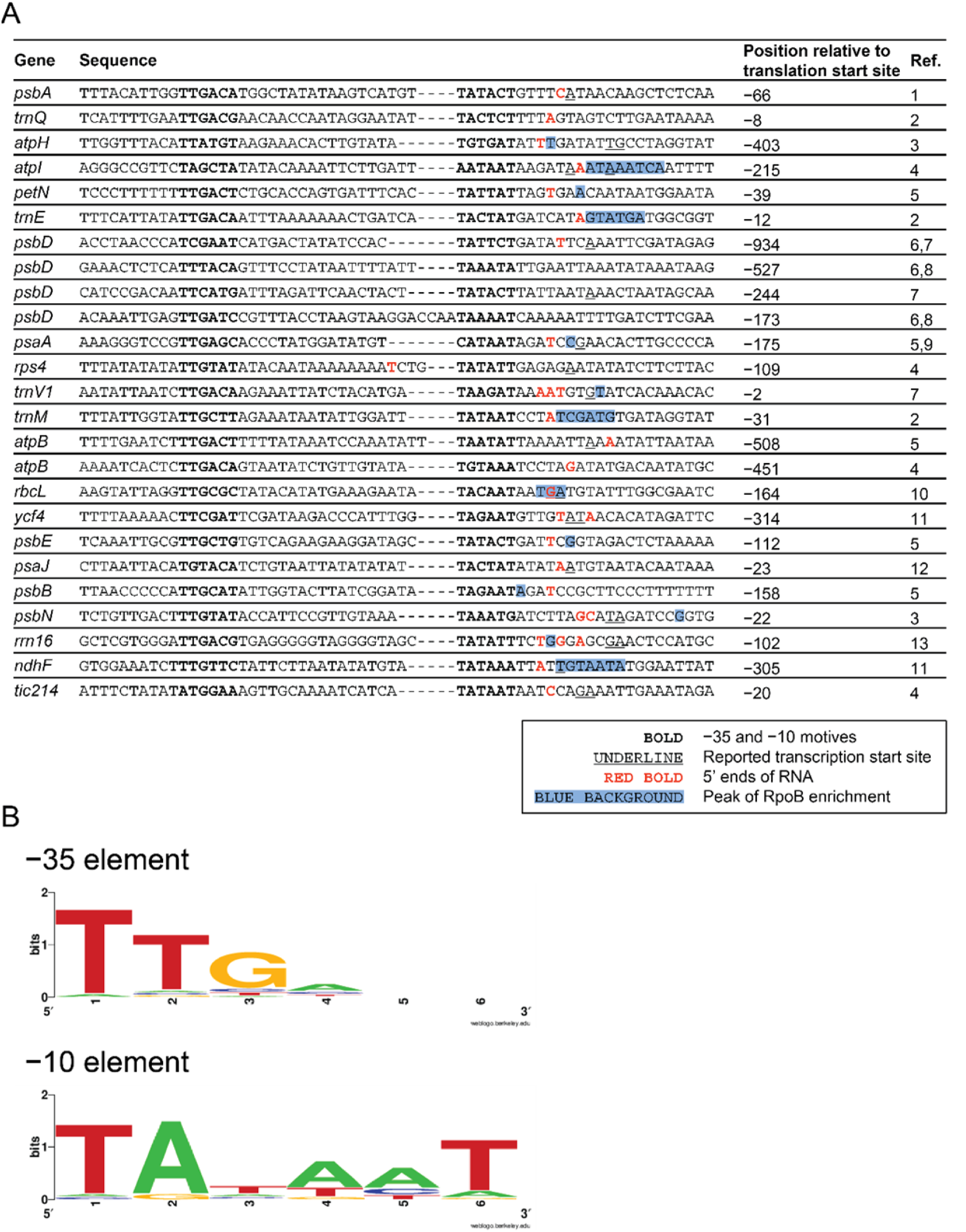
PEP promoters identified in Arabidopsis. A. Alignment of PEP promoters. Positions of the last nucleotides (20 nt from the −10 element) relative to translation start sites are indicated. Transcription start sites reported previously and identified with TSS RNA-seq in this study are represented with underlines and red colors, respectively. Blue background indicates the position with highest RpoB enrichment downstream of each −10 element. ^1^Liere et al., 1995; ^2^Kanamaru et al., 2001; ^3^Zghidi et al., 2007; ^4^Swiatecka-Hagenbruch et al., 2007; ^5^Ishizaki et al., 2005; ^6^Hoffer and Christopher, 1997; ^7^Hanaoka et al., 2003; ^8^Shimmura et al., 2008; ^9^Fey et al., 2005; ^10^Hakimi et al., 2000; ^11^Favory et al., 2005; ^12^Nagashima et al., 2004; ^13^Sriraman et al., 1998. B. Sequence logos for −35 and −10 elements of PEP promoters listed in A. Images were generated by using WebLogo (https://weblogo.berkeley.edu/logo.cgi) with default settings.

**Figure S6.**
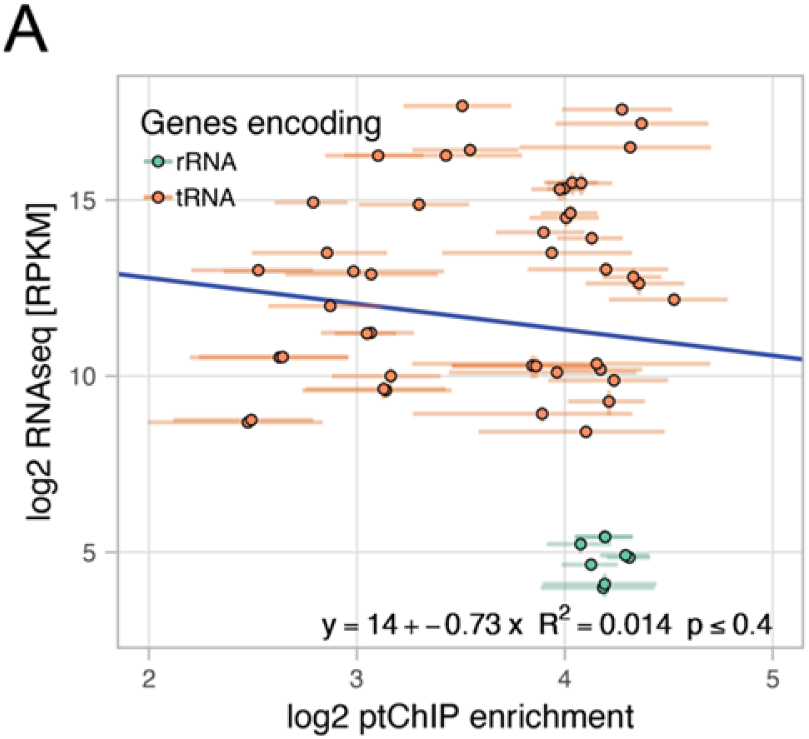
Correlation of PEP binding with steady state levels of RNA. A. No correlation on rRNA and tRNA genes between RpoB ptChIP-seq enrichments and total RNA-seq signals. Enrichment levels of RpoB ptChIP-seq in 14 days old Col-0 wild type plants (this study) and RPKM normalized total RNA-seq signals from a similar developmental stage (Thieffry et al., 2020) were compared on annotated rRNA and tRNA genes. Data points are color-coded by function of the corresponding genes and show averages from three independent replicates. Error bars indicate standard deviations. Blue line represents the linear regression model.

**Table S1.**
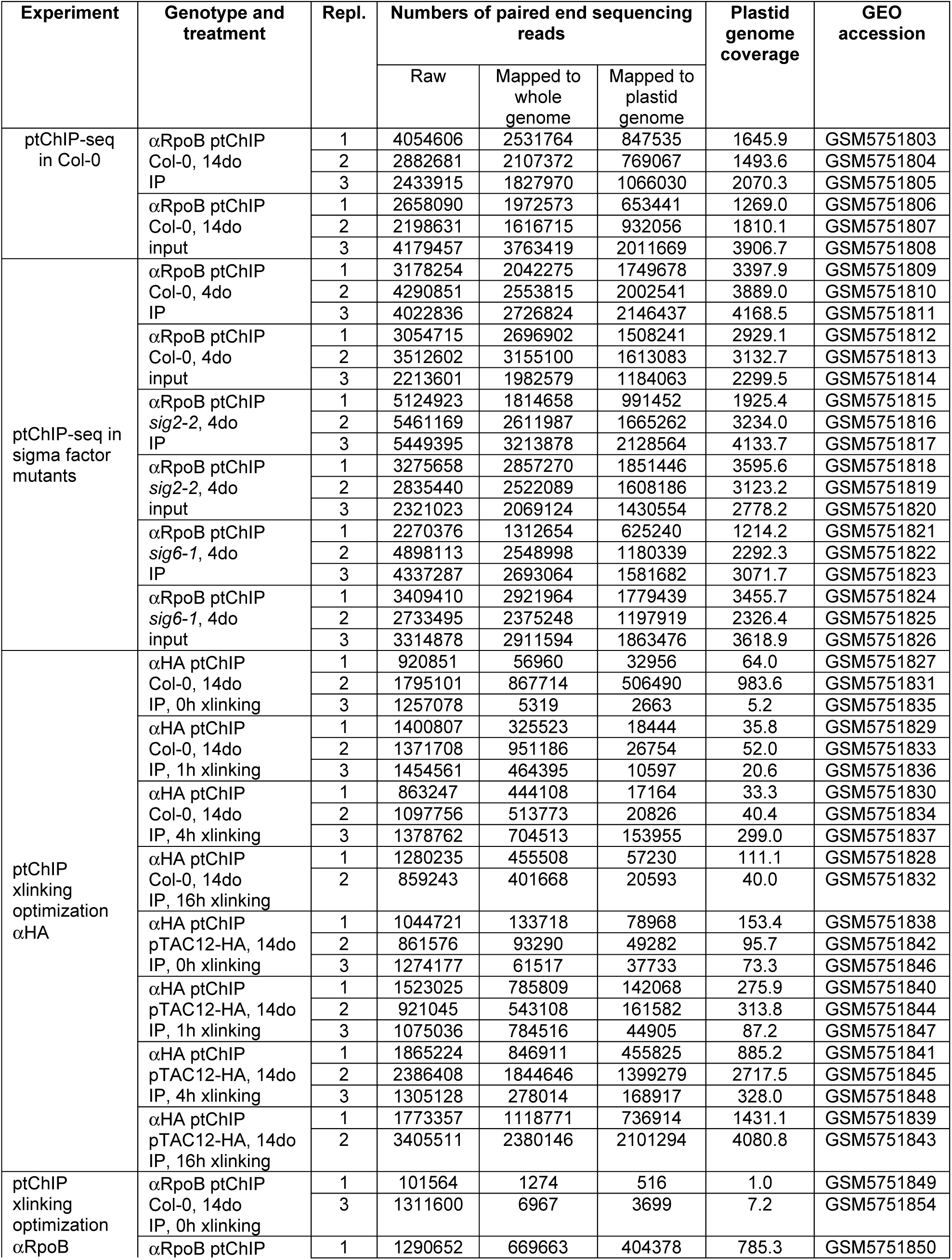
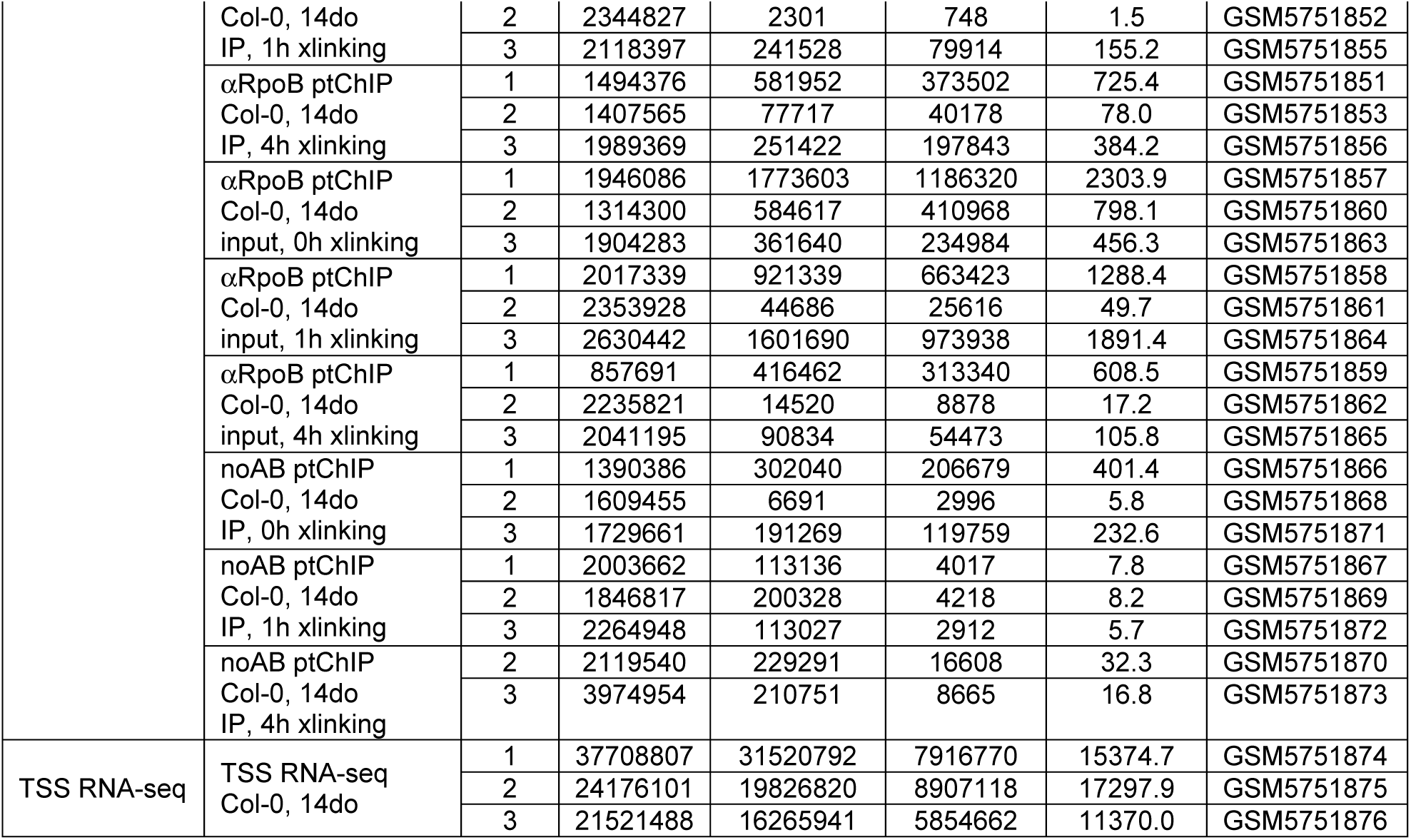
High throughput sequencing datasets generated in this study.

**Table S2.**
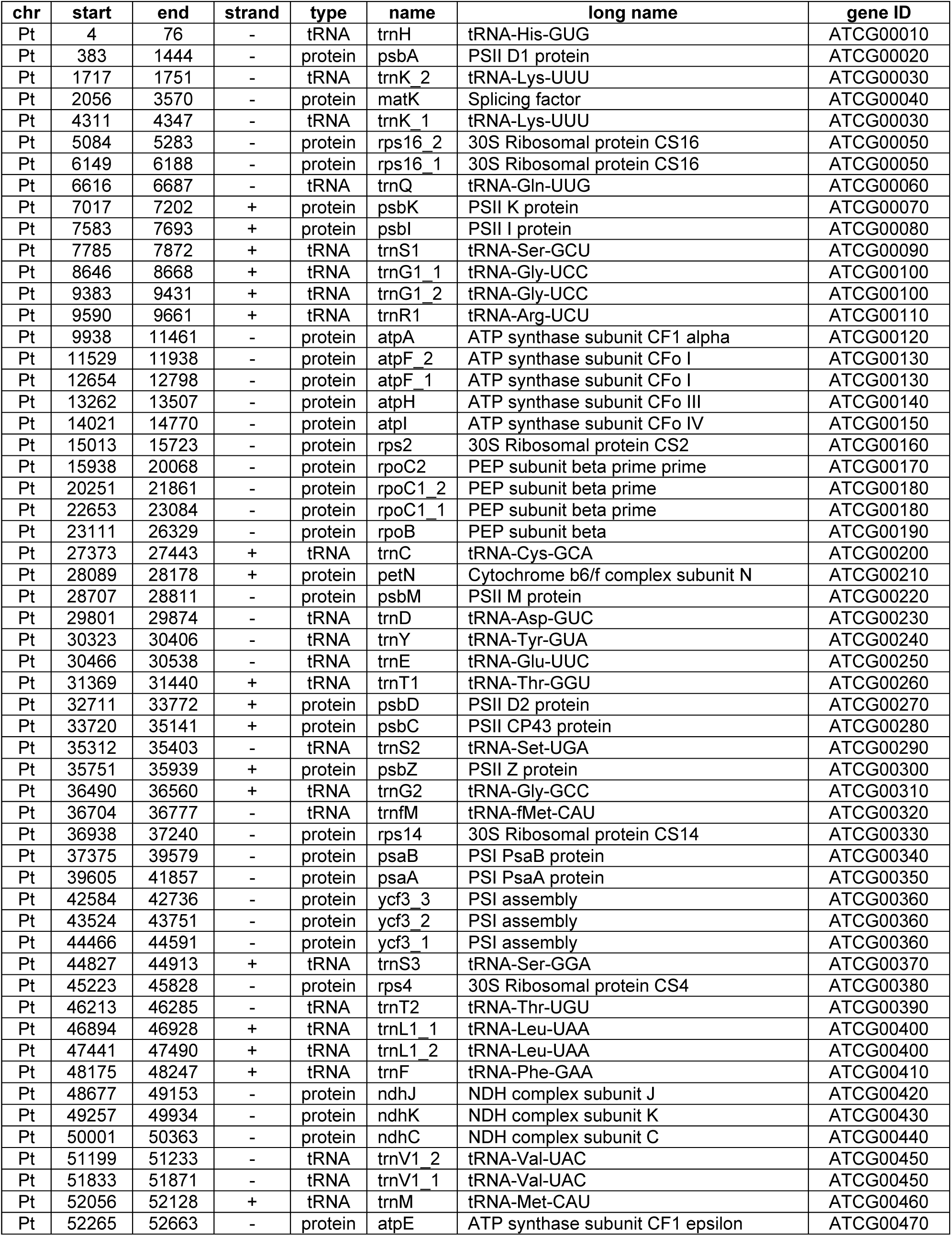
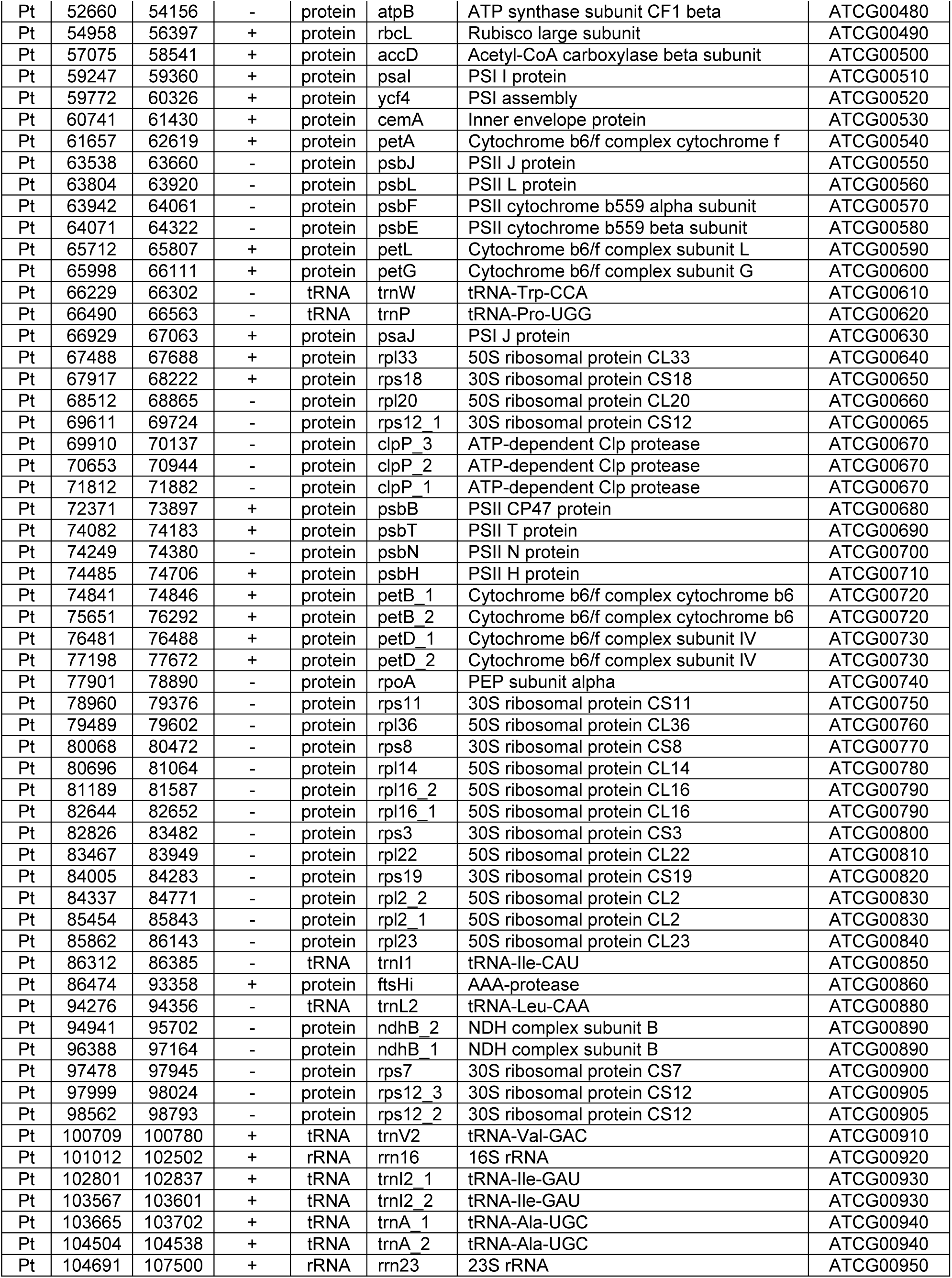
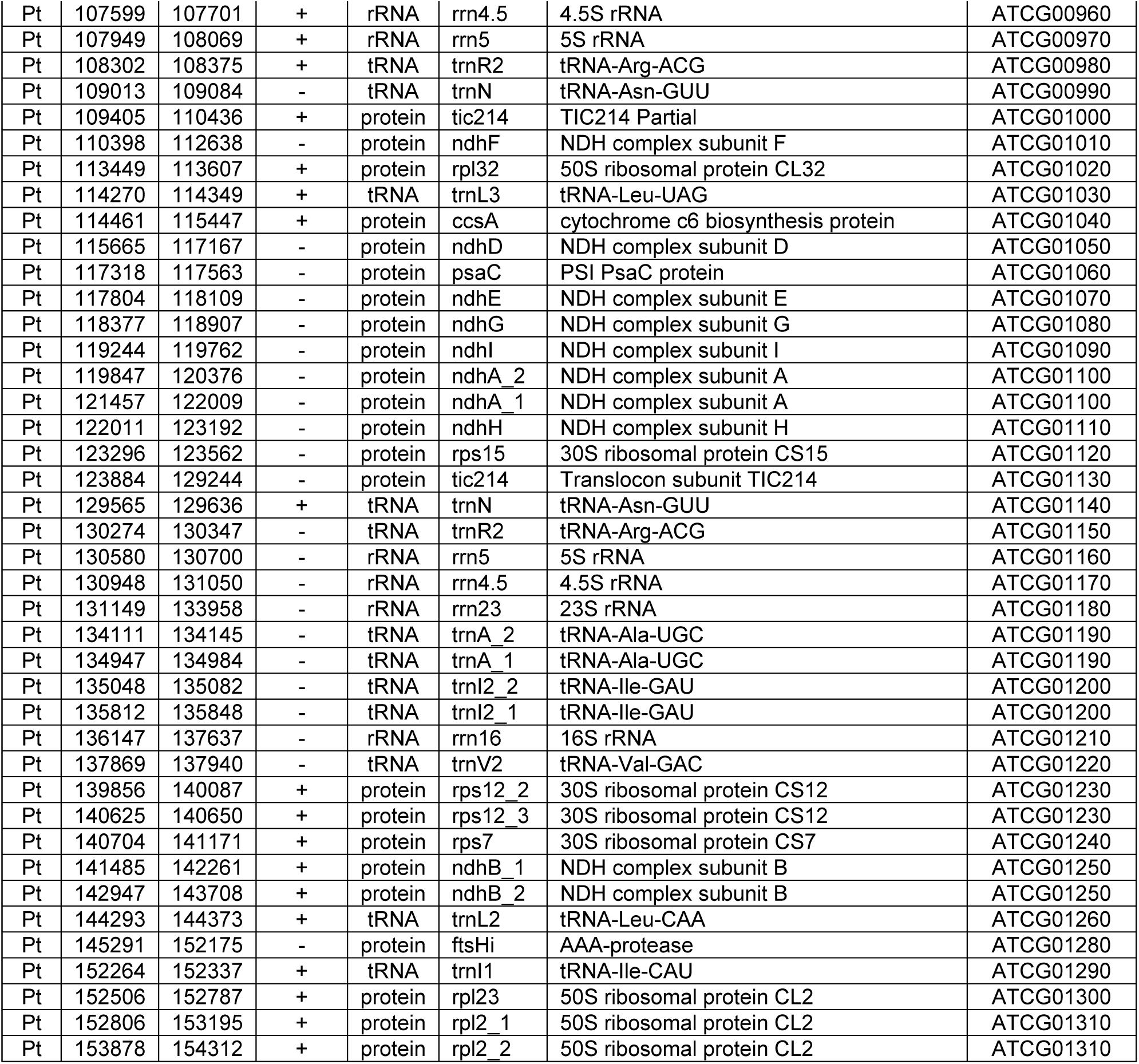
Annotation of plastid-encoded genes in Arabidopsis used in this study.

